# Whole genome sequencing evidences higher rates of relapse in *Clostridioides difficile* infection caused by PCR ribotype 106

**DOI:** 10.1101/2021.03.23.436720

**Authors:** Loreto Suárez-Bode, Carla López-Causapé, Antonio Oliver, Ana Mena

## Abstract

Increasing prevalence and widespread of *Clostridioides difficile* infection (CDI) caused by the epidemic DH/NAP11/106/ST-42 has been observed worldwide, probably fostered by its great capacity to produce spores or the higher resistance rates found in some strains. Previous studies have also attributed higher recurrence rates to RT106 as compared to others. This study describes the genetic analysis by whole genome sequencing (WGS) of primary and recurrent isolates of RT106 to determine if the higher rate of recurrence associated to RT106 is due to relapses, caused by the same strain, or reinfections, caused by different strains. The whole-genome sequences of ten primaries and fourteen recurrent RT106 isolates from ten patients were obtained to determine MLST profiles, resistance mutations and phylogenetic relatedness between the different isolates by comparative single nucleotide variants (SNV) analysis using the *C. difficile* DH/NAP11/106/ST-42 genome as reference (GenBank accession: GCA_002234355.1). All isolates were classified as ST42 and SNVs comparative analysis showed that those belonging to the same patient were isogenic (differed by ≤3 SNVs), with one exception (6 SNVs); while strains belonging to different patients were not (differing by 4 to 43 SNVs) with two exceptions of isogenic isolates from different patients, suggesting putative transmission events. Phylogenetic analysis suggested the presence of similar local epidemic lineages, except for one patient whose isolates clustered with different US isolates. All the isolates, except those clustering with the US lineage, were also associated to moxifloxacin resistant since all of them were resistant by agar diffusion (MIC >32) and contained the Thr82Ile mutation in *gyrA*. Our results evidence that recurrent *Clostridioides difficile* infections caused by RT06/ST42 are mainly due to relapses, caused by primary strains, showing the great capacity of RT106/ST42 to persist and cause recurrences as compared to other ribotypes.

## INTRODUCTION

The increasing prevalence and widespread of the epidemic DH/NAP11/106/ST-42 has been reported in the last years in regions where it was previously rarely found. Although initially identified in the United Kingdom in 1999, RT106, is now one of the predominant strains in Europe, and it is also becoming the most prevalent strain in the United States where RT106 has replaced RT027 as the most prevalent ribotype recovered from community-associated *C. difficile* infections (CDI) and where has also been reported as the second most prevalent molecular type in acute-care hospitals [1–5].

The last National surveillance study performed in 2013 in Spain suggested ribotypes 078/126, 014 and 001/072 as the most prevalent in this country [6] that was in accordance with our local surveillance data where ribotypes 014 and 078 were described as the most prevalent [7]. However, later studies showed changes in the epidemiology of *C. difficile* in Spain, such as the 2015-2016 surveillance analysis published by Suárez-Bode *et* al. [8] revealing RT106 as the most prevalent ribotype both in community associated (CA)- and health-care facility associated (HCFA-) CDIs. High prevalence has also been reported for RT106 in another Spanish hospital where it was also associated to large outbreaks and nosocomial transmission even between non-related patients [9].

The growing incidence and geographic distribution of the epidemic RT106 seems to be favored by different features that some authors have described to be associated with these strains as high level of sporulation, combined with moderate toxin production, and its resistance to environmental decontamination, as compared with other ribotypes and reference strains [10, 11]. On the other hand, antimicrobial resistance may be an important associated factor since it is a major contributor to the pathogenesis and global dissemination of epidemic strains, like BI/NAP1/ 027 [12–14] and, in this sense, highest rates of resistance have also been related to RT106 strains among adults in Ireland, Scotland or Spain, specially associated to fluoroquinolone resistance [8, 15, 16]. Moreover, RT106 has also ten accessory genomic elements (AGEs) with unique sequences that could be related with intestinal mucosal adhesion, biofilm formation, and sporulation although the function of the proteins transcribed from these sequences need to be further studied [1, 17].

It has been speculated that the ability of RT106 to produce high levels of spores may favor to cause hospital outbreaks and recurrent disease more effectively than other ribotypes [1, 18]. However, more data is needed to demonstrate these beliefs. Different studies have revealed a higher recurrence rate exhibited by RT106 strains in comparison with other major ribotypes [3, 8]. In fact, Kociolek *et* al. identified several genes strongly associated with strain DH/NAP11/106 that could play a role providing bacterial competitive advantages and association with recurrent CDI although further investigation is needed [17].

Whole-genome sequencing (WGS) is the preferred method for tracking *C. difficile* due to its exceptional sensitivity in discriminating strain genetic relatedness and identifying putative transmission events [19]. Single nucleotide variants (SNV)-based analysis has been widely adopted for CDI surveillance and has revealed some new evidence about transmission dynamics and recurrent infections [20]. The main objective of the present study was to analyze by WGS primary and recurrent RT106 isolates from patients with recurrent CDI in order to prove if the higher rate of recurrence associated to this ribotype is in fact due to relapses, caused by the same strain, or to reinfections, caused by different strains.

## METHODS

### Patients selection and *C. difficile* isolation

The study was conducted at the reference Hospital of the Balearic Islands (Spain) a 700-bed tertiary-care university hospital providing medical care for approximately 750,000 inhabitants. A cohort of patients previously diagnosed of recurrent episodes of CDI caused by RT106 between years 2016 and 2018 were selected and included in the study. Recurrence was considered as a new episode of CDI during the 8 weeks following the previous resolved episode but also, “late recurrences” (more than 8 weeks following the previous resolved episode) were included, as previously described [8].

Specimens were cultured on selective cycloserine–cefoxitin–fructose agar plates (CLO agar; bioMérieux, Marcy l’Etoile, France) in an anaerobic chamber and incubated at 37°C for 48 h, as previously described [8]. All the isolates were obtained from stool specimens from hospital (inpatients), ambulatory and primary care (outpatients) adult patients and children below two with confirmed diagnosis of CDI. *C. difficile* isolates were frozen and stored at −80°C to be processed for WGS on a later stage. Isolates were named as “CD” with the patient number and “a” letter for initial episodes (first episode), “b” for the first recurrent episode (second episode), “c” for the second recurrent episode (third episode).

Initial episodes of CDI were classified as health-care facility associated (HCFA-CDI) (hospital or community onset), community associated (CA-CDI) or as indeterminate disease according to the established definitions [21].

### Molecular characterization of the isolates

The selection of isolates from patients with recurrent CDI caused by RT106 was carried out through the previous characterization of isolates by high-resolution capillary gel-based electrophoresis PCR-ribotyping using the protocol previously described by Fawley et al. [22]. Electrophoerograms were obtained by Gene Mapper (v.4.0.) and analyzed by the Webribo database [23].

### Whole-genome sequencing and analysis

All the included isolates underwent whole-genome sequencing. Genomic DNA was isolated using a commercial extraction kit (High Pure PCR template preparation kit; Roche Diagnostics). Indexed paired-end libraries were preparedwith the Nextera XT DNA library preparation kit (Illumina Inc, USA) and then sequenced on an Illumina MiSeq^®^ benchtop sequencer with MiSeq reagent kit v3 (Illumina Inc., USA), resulting in 300 bp paired-end reads. MLST analysis was performed by using the online tool MLST-2.0. (https://cge.cbs.dtu.dk/services/cge/) [24].

To perform pairwise single-nucleotide variant (SNV) analysis, paired-ended reads were aligned to the *C. difficile* DH/NAP11/106/ST-42 genome (GenBank accession: GCA_002234355.1) using Bowtie 2 v2.2.4 (http://bowtie-bio.sourceforge.net/bowtie2/index.shtml) and, eventually, pileup and raw files were obtained by using SAMtools v0.1.16 (https://sourceforge.net/projects/samtools/files/samtools/) and PicardTools v1.140 (https://github.com/broadinstitute/picard). The Genome Analysis Toolkit (GATK) v3.4-46 (https://www.broadinstitute.org/gatk/) was used for realignment around InDels. SNPs were extracted from the raw files if they met the following criteria: a quality score (Phred-scaled probability of the sample reads being homozygous reference) of at least 50, a root-meansquare (RMS) mapping quality of at least 25 and a coverage depth of at least three reads, excluding all ambiguous variants. MicroInDels were extracted from the total pileup files applying the following criteria: a quality score of at least 500, an RMS mapping quality of at least 25 and support of at least one-fifth of the covering reads.

As no definitive criteria have been established for relatedness definition, and based on previous works, we considered isolates collected <124 days apart as isogenic if they differed by ≤2 single nucleotide variants, and those collected between 124-364 days apart as isogenic if they differed by ≤3 single nucleotide variants. Unrelated isolates were considered those differing ≥ 10 SNVs [19, 20]. SNV calling is also a very useful method for local transmission analysis, therefore we considered plausible transmission events between patients when isolates differed ≤2 SNPs within ≤90 days.

In order to analyze the relatedness among isolates, genomes were *de novo* assembled using SPAdes and a core genome phylogenetic reconstruction was built with Parsnp from the Harvest Suite package v1.2 with default parameters forcing the inclusion of all genomes and randomly selecting the reference genome (flags: -c/-r!) and including other available ST42 genomes (ENA Bioproject number: PRJNA340238) [17].

The nucleotide sequences for the twenty-four genomes included in this study have been deposited at DDBJ/ENA/GenBank under the following number project: PRJEB43620.

### Antimicrobial susceptibility and mechanisms of resistance analysis

Susceptibility to metronidazole, vancomycin, erythromycin and moxifloxacin using gradient antibiotic strips (Etest^®^, BioMérieux, Marcy l’Etoile, France) on Brucella agar plates (BD Biosciences, Oxford, UK) was tested for all the isolates. MIC breakpoints used were those established by the European Committee on Antimicrobial Susceptibility Testing (EUCAST) for metronidazole (>2 mg/L) and vancomycin (>2 mg/L) and ECOFF value for moxifloxacin (4 mg/L) and erythromycin (2 mg/L) (v.8.0; (http://www.eucast.org/clinical_breakpoints/).

Draft genomes were screened to identify acquired genes mediating antimicrobial resistance or chromosomal mutations of resistance for *murG, rpoB, gyrA, gyrB, cfr, tetM, catP* or *ermB* genes and also by using the online tool Resfinder (https://cge.cbs.dtu.dk/services/ResFinder/) [25, 26].

## RESULTS

### Patients and isolates characteristics

Ten patients with recurrent diagnosis of CDI caused by RT106 were finally included in the study, with a total of twenty-four isolates, ten from initial infections and fourteen from recurrent episodes (ten of them from first recurrent episodes and the remaining four from the second recurrent episodes).

The mean age of the patients was 64 (2-92y) years old and the mean time between different recurrent episodes was 24.5 (19-127) days, including twelve recurrent episodes, according to the strict definition, and two “late recurrences” (78 and 127 days after the previous episode). Primary CDI episodes were considered, three as HCFA-CDI, six as CA-CDI and one case was classified as an indeterminate disease. Table 1 summarizes the main epidemiological data of patients and their isolates.

**Table 1.** Epidemiological data of patients and isolates including MLST profile and pairwise single-nucleotide variant (SNV) analysis.

| Patients | Patients age | Gender | Isolates | Days between episodes | Type of episode | CDI associated adquisition | PCR-RT/MLST | SNVs withing preceding isolate |
| --- | --- | --- | --- | --- | --- | --- | --- | --- |
| **CD1** | 83 | F | CD1a | 0 | Primary | HCFA-ICD | R106/ST42 |  |
|  |  |  | CD1b | 23 | First recurrence |  | R106/ST42 | 2 |
|  |  |  | CD1c | 20 | Second recurrence |  | R106/ST42 | 0 |
| **CD2** | 58 | F | CD2a | 0 | Primary | HCFA-ICD | R106/ST42 |  |
|  |  |  | CD2b | 127 | Late recurrence |  | R106/ST42 | 1 |
| **CD3** | 34 | M | CD3a | 0 | Primary | Indeterminate | R106/ST42 |  |
|  |  |  | CD3b | 78 | Late recurrence |  | R106/ST42 | 0 |
| **CD4** | 61 | M | CD4a | 0 | Primary | CA-ICD | R106/ST42 |  |
|  |  |  | CD4b | 24 | First recurrence |  | R106/ST42 | 3 |
| **CD5** | 92 | F | CD5a | 0 | Primary | CA-ICD | R106/ST42 |  |
|  |  |  | CD5b | 24 | First recurrence |  | R106/ST42 | 0 |
| **CD6** | 87 | F | CD6a | 0 | Primary | HCFA-ICD | R106/ST42 |  |
|  |  |  | CD6b | 24 | First recurrence |  | R106/ST42 | 0 |
|  |  |  | CD6c | 25 | Second recurrence |  | R106/ST42 | 0 |
| **CD7** | 2 | M | CD7a | 0 | Primary | CA-ICD | R106/ST42 |  |
|  |  |  | CD7b | 24 | First recurrence |  | R106/ST42 | 1 |
| **CD8** | 38 | F | CD8a | 0 | Primary | CA-ICD | R106/ST42 |  |
|  |  |  | CD8b | 19 | First recurrence |  | R106/ST42 | 0 |
| **CD9** | 69 | M | CD9a | 0 | Primary | CA-ICD | R106/ST42 |  |
|  |  |  | CD9b | 32 | First recurrence |  | R106/ST42 | 0 |
|  |  |  | CD9c | 43 | Second recurrence |  | R106/ST42 | 0 |
| **CD10** | 64 | M | CD10a | 0 | Primary | CA-ICD | R106/ST42 |  |
|  |  |  | CD10b | 43 | First recurrence |  | R106/ST42 | 6 |
|  |  |  | CD10c | 34 | Second recurrence |  | R106/ST42 | 3 |

### Whole-genome sequencing analysis

All the isolates included in the study were classified as sequence-type (ST) 42 based on the MLST analysis, as was expected for ribotype 106.

SNVs comparative analysis (variant calling) of isolates belonging to the same patient showed that recurrent episodes of patients CD1, CD2, CD3, CD5, CD6, CD7, CD8 and CD9 were caused by isogenic strains since isolates from the same patient differed by 0 to 2 SNPs and were collected less than 124 days apart. Furthermore, variant calling analysis showed only 3 SNVs of difference between isolates belonging to patient CD4 and also between isolates CD10a and CD10b with isolate CD10c, all of them collected less than 124 days apart and, therefore, not meeting the strict definition of isogenic strains. However, we consider these strains as “presumably isogenic” isolates. Finally, isolates CD10a and CD10b differed by 6 SNPs being, therefore, non-isogenic but not unrelated strains (Figure 1).

**Figure 1.**
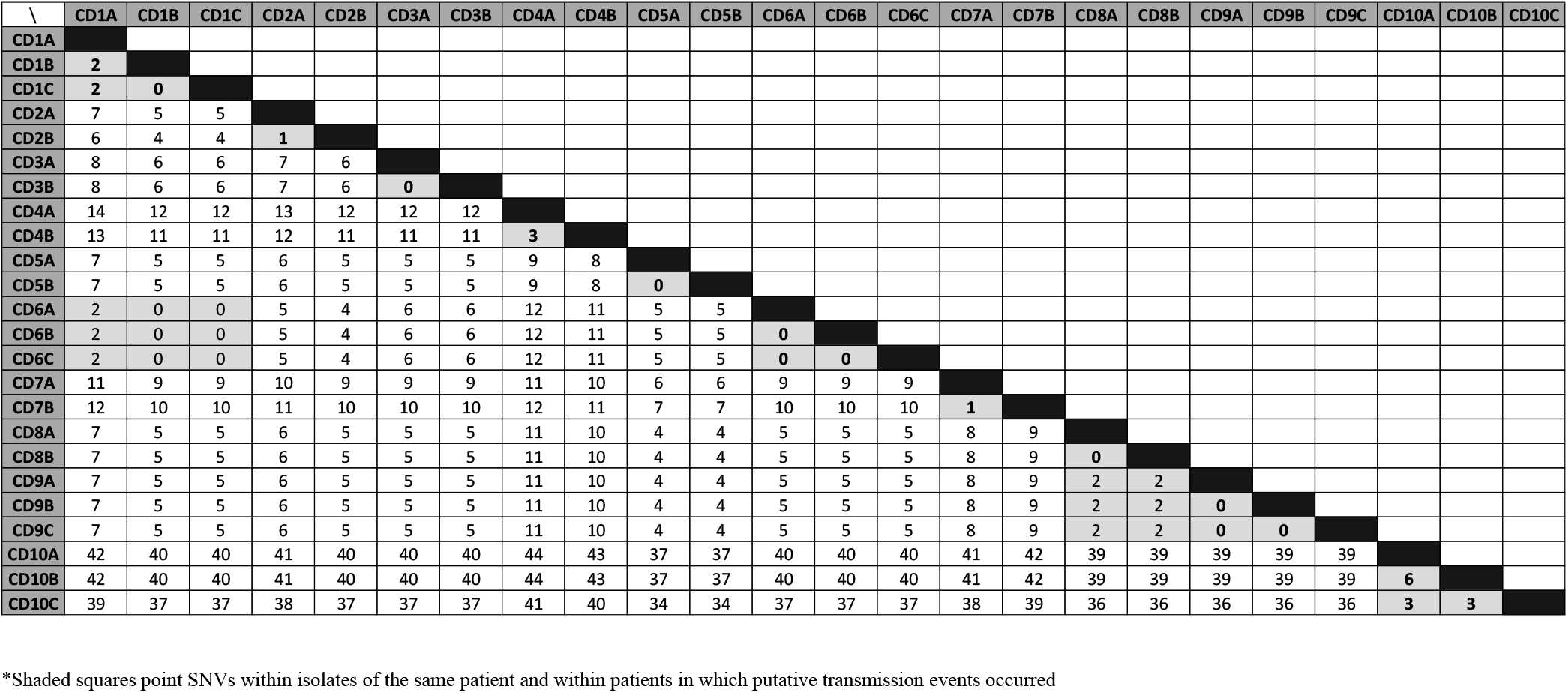
SNP comparative analysis within the different patient isolates.

Overall, these results showed that all the recurrent episodes were due to isogenic strains with the initial ones, being able to classify all these cases as relapses, with the exception of the first recurrent episode of patient CD10, where a related but non isogenic strain was found, and, therefore, not concluding if a relapse or a reinfection occurred in that case.

Isolates belonging to different patients differed by 4 to 43 SNPs with the exception, of the isolates of patients CD1 and CD6 on one side, and isolates of patients CD8 and CD9 on the other, differing by only 2 SNPs, and suggesting, both cases, a plausible transmission events within these patients. Patients CD1 and CD6 were admitted to the same hospital at the same time and CDI cases occurred within ≤90 days. On the other hand, patients CD8 and CD9 were also admitted in the same hospital but not during the same period, however CDI episodes occurred within ≤90 days (Figure 2).

**Figure 2.**
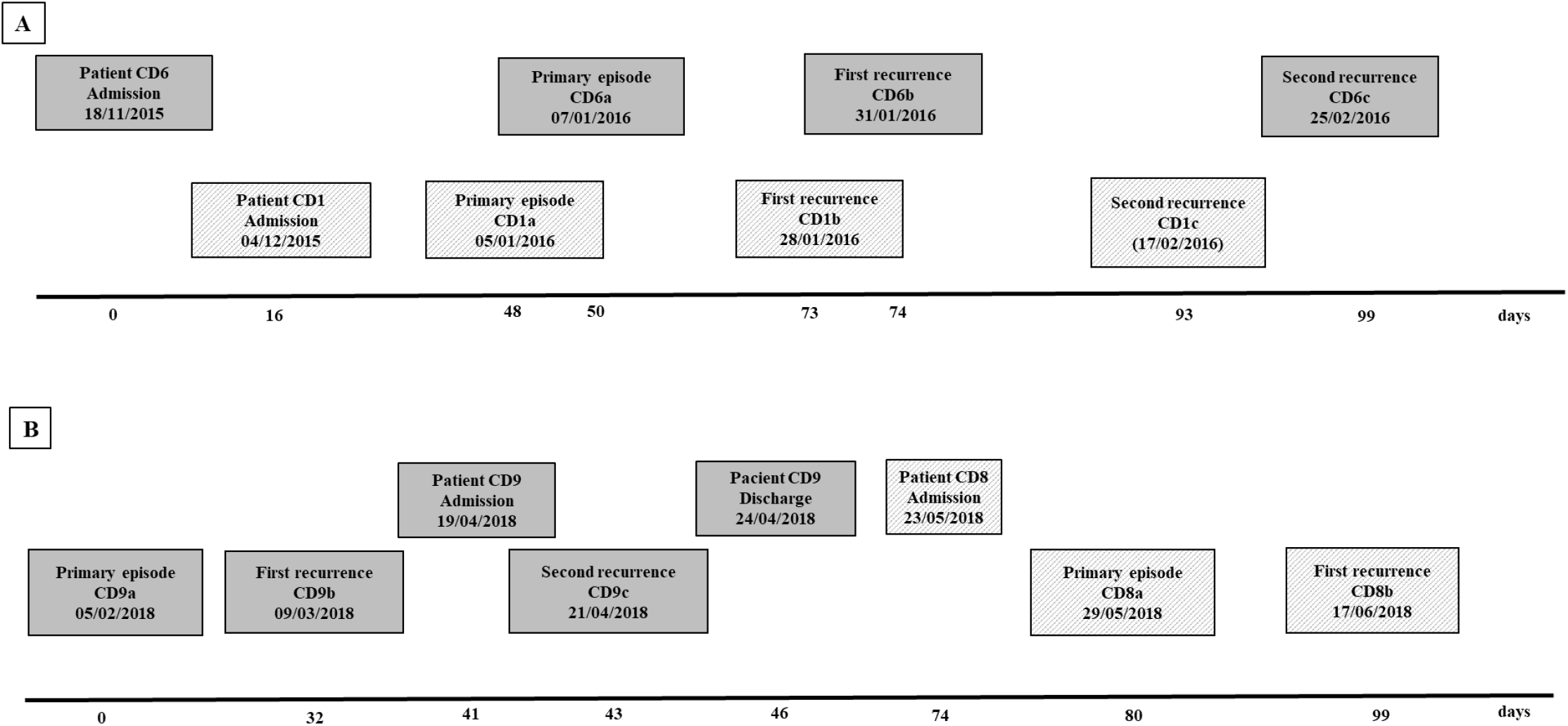
Representation of the two plausible transmission events within patients a) CD1 and CD6, admitted during a common period; and b) patients CD8 and CD9, not admitted at the same time at hospital.

Comparative analysis between isolates from the same patients showed only two patients with nonsynonymous changes between their primary and recurrent isolates (Table 2), one of them (Ala66Val) located at the S-layer protein SlpA. Therefore, no common mutations were found either between recurrent isolates.

**Table 2.** Nonsynonymous changes found in recurrent isolates with respect to the primary ones.

| Isolate | Position | Reference allele | Isolate allele | Gene | Protein | Amino acid change |
| --- | --- | --- | --- | --- | --- | --- |
| CD7b | 3118341 | G | A | *slpA* | S-layer protein SlpA | Ala166Val |
| CD10b | 1645698 | G | T | CGC51_RS07795 | response regulator transcription factor | His162Asn |
| CD10b | 1986896 | G | T | CGC51_RS09380 | 3-dehydroquinate synthase | Ala117Ser |
| CD10b | 2458725 | C | A | CGC51_RS11610 | cell division protein FtsK | Gly420* |

### Phylogenetic analysis

Core phylogenetic analysis showed three main branches grouping all the isolates in five clusters. Isolates from patient CD10 cluster with some US isolates, included in the phylogenetic tree for comparison, showing greater relatedness with them while isolates from the rest of patients (CD1-CD9) gathered in four clusters showing greater similarity between them. Moreover, this analysis supported the direct transmission between patients CD1 and CD6 and patients CD8 and CD9 clustering their respective isolates together in both cases (Figure 3). Synonymous and nonsynonymous changes found in the isolates regarding the reference strain also confirm the observed relatedness in the phylogenetic tree since common changes were found in all the isolates for patients CD1 to CD9, differing from those observed in patient CD10 isolates, confirming the presence of a different lineage in this case. All mutations and nucleotid changes found for all the isolates are summarized in the Supplementary material (see web-only Supplementary Table S1).

**Figure 3.**
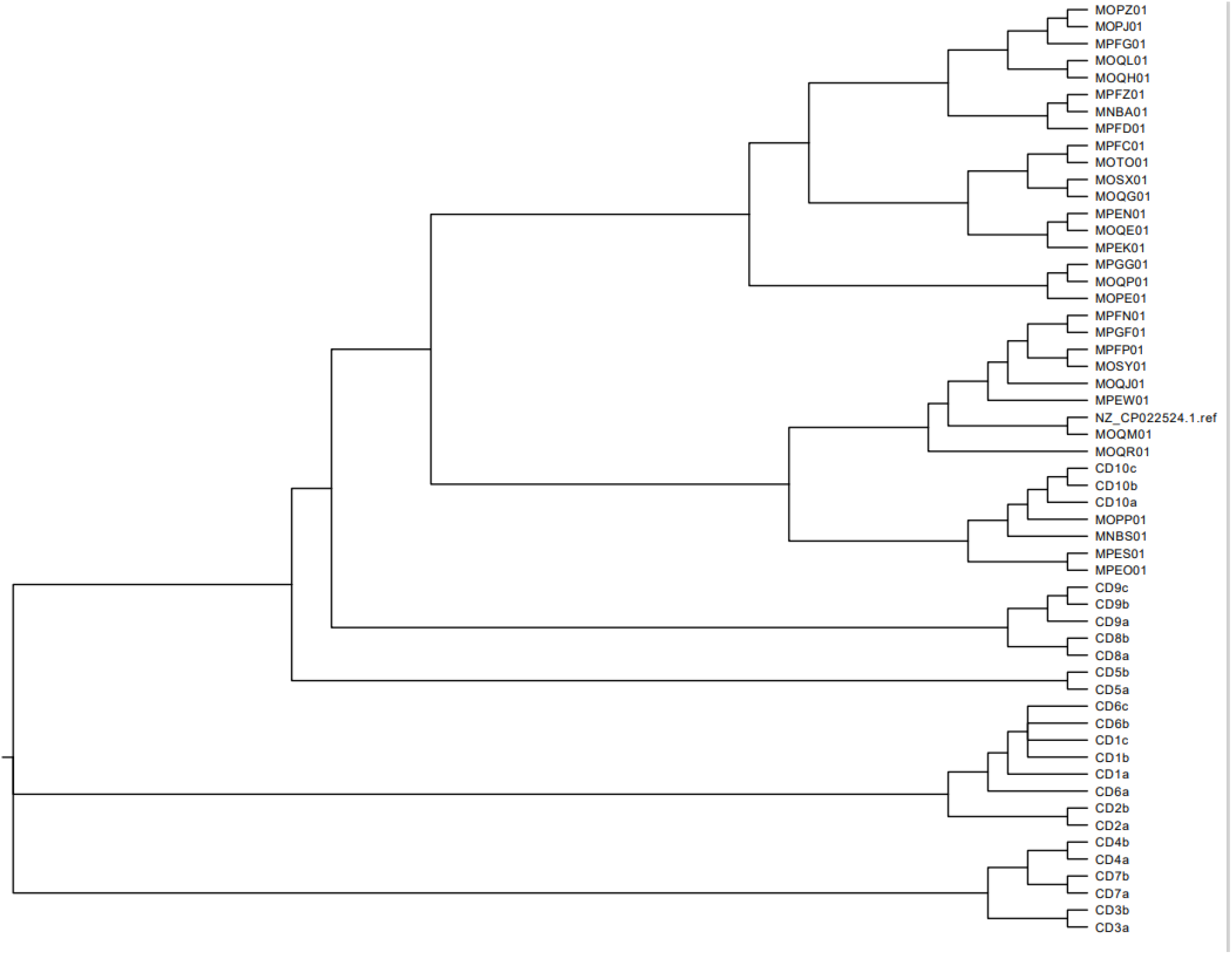
Phylogenetic tree showing representatives of the primary and recurrent isolates from the ten studied patients (CD1-CD10) and other 30 available ST-42 isolates included for comparison.

### Antimicrobial susceptibility and mechanisms of resistance

All isolates were found to be susceptible to metronidazole and vancomycin and resistant to moxifloxacin with MICs > 32 mg/L, except for isolates of patient CD10 that were also susceptible to moxifloxacin. Only isolate CD3b was considered resistant to erythromycin (MIC 2 mg/L). WGS analysis showed the Thr82Ile mutation in *gyrA* for moxifloxacin resistant isolates with no other resistance mutations for the screened genes and no other acquired resistant genes were found with the Resfinder analysis.

## DISCUSSION

There are several factors that seems to contribute to the emergence and widespread of RT106 strains as the high level of sporulation demonstrated for this ribotype or higher rates of antibiotic resistance reported by different studies [8,10,18]. Moreover, some of these advantages seems to be related with the ability of RT106 to produce recurrent disease more effectively than other ribotypes. Kociolek *et* al. observed that CDI caused by DH/NAP11/106 was more likely to result in multiple CDI relapses (40% vs 8%; *P*=.05) when they studied the molecular epidemiology by restriction endonuclease analysis (REA) groups among children with multiple CDIs [3]. Based on the higher recurrence rate (26%) that was also observed for RT106 in our previous study as compared to other major ribotypes (*p*<0.05), we sequenced all the isolates from ten different patients with recurrent CDI caused by this ribotype to find out if recurrences were caused by the same strain, considering them as relapses, or by different strains of the same ribotype, causing reinfections. Variant calling analysis evidenced that recurrent episodes were caused mostly by clonal strains since recurrent isolates were isogenic (≤2 SNPs within isolates) with the strain responsible of the previous episode, with the exception of one patient where a recurrent isolate was non isogenic but closely related with the previous strain (only 6 SNPs within isolates). Based on these findings, it seems that recurrent *C. difficile* infections caused by RT06/ST-42 are due to relapses caused by a clonal strain, suggesting a higher capacity of these strains to persist and cause greater number of recurrences than other ribotypes. As other authors have commented previously, we cannot assume that relapses are caused by the persistence of strains in the gut since spores of *C. difficile* are capable to persist for a long time in the environment being unable to rule out infections caused by the same strain [9]. However, the fact that we did not observe isogenic strains among the majority of patients (differing between 4 and 43 SNPs) makes this theory less likely, since it would be expected to find more isogenic strains among the different patients if they had been reinfected with persistent spores of the same strain present in the environment. In these sense, a new important study has recently demonstrated a novel mechanism employed by *C. difficile* spores to gain entry into the intestinal mucosa via pathways dependent on host fibronectin-α5β1 and vitronectin-αvβ1 where the exosporium protein BclA3, on the spore surface, is required for both entry pathways contributing to the recurrence of disease [27].

No common mutations were detected in the recurrent isolates to explain adaptation mechanisms or some kind of advantage to persist in the human gut for these isolates, nevertheless, analysis of accessory genome elements (AGEs) need further study since some mechanisms related with adaptation or persistence could be related with them. Many of the CDSs identified in the genome of *C. difficile* are associated with adaptation and proliferation in the gastrointestinal tract (germination, adhesion, and growth) and survival in challenging suboptimal environments (endospore formation) supporting the view that *C. difficile* lives within a highly dynamic niche and is able to spend a long time coexisting with its host [28]. In fact, comparative genomic studies found that RT106 had the lowest conservation of the core genes present in the reference strain CD630 of all ribotypes tested, while showed 100% conservation of the divergent sequences present in the hipervirulent RT027, which may represent genes associated with increased virulence [1], although more studies are needed to associate gene sequences with virulent phenotypes and/or epidemic strains. A recent study published by Roxas *et* al., shows clade-specific properties, including those conferred by genes within the genomic island GI1 that could explain the higher emergence of this clones by assessing virulence-associated phenotypes including motility, toxin production, biofilm production or adhesion to collagen [5]. However, there must be other factors that play a role as well and more studies are needed to confirm and better understand these process. Moreover, two plausible transmission events between patients were also documented since their isolates were isogenic and episodes took place during the same period of time (within ≤90 days). In one of the cases, the two patients involved were admitted to the same hospital at the same time while on the other case patients were at the same hospital but at different times. García-Fernández *et* al., showed a frequent within-hospital transmission of healthcare-associated ribotypes, including ribotype 106, and they also observed that a significant proportion of transmissions occurred either indirectly, through environmental contamination with *C. difficile* spores, or from reservoirs outside of CDI patients, such as asymptomatically colonized patients or staff [9].

Phylogenetic analysis showed a close relatedness between isolates from all the patients suggesting the presence of similar local epidemic linages, except in one patient in which his isolates clustered with other US isolates. Beside this, all the isolates belonging to the local epidemic lineages were resistant to moxifloxacin unlike the other US lineage, showing the Thr82Ile mutation in *gyrA* previously described. Resistance to fluoroquinolones in DH/NAP11/106 strains have been previously reported in Europe among adults in different countries like Ireland, Scotland or Spain [8, 15, 16], in contrast to isolates reported from North America where resistance to moxifloxacin has not been associated to this ribotype [17].

Our work highlights the importance of WGS to determine relapses, caused by the same strain, or reinfections, caused by different strains. Current definitions are based only on the lapse since previous episode (8 or less weeks), but there are “late” recurrences (more than 8 weeks) that might be caused by the same strain, highlighting the need to change the standard definition of relapse. Additionally, as other authors have considered, to differentiate between relapses and reinfections might be important for controlling CDI, either through interventions to manage *C. difficile* transmission, or implementing treatment policies requiring a different handling or even individualized therapeutic strategies [29].

Our study has some limitations since has been performed in a single center with a limited number of patients, however, it is significant that all the recurrent isolates except one, very closely related with the previous (6 SNPs), were isogenic with respect to their primary isolates. Moreover, despite being logical thinking that there is possible limitation identifying relapses or transmission events using WGS of a single colony, Balaji *et* al. [30], identified rare within-host genetic diversity of *C. difficile* in stool of children with CDI or asymptomatic carriage, suggesting that WGS of a single colony from stool will appropriately characterize isolate clonality and putative transmission events in the majority of cases.

In conclusion, we have evidenced by WGS higher rates of relapse in CDI caused by RT106/ST42 strains, mainly represented by a local epidemic strains associated to fluoroquinolone resistance. However, additional studies are needed to better understand the mechanisms of persistence in these strains to cause recurrent CDI.

## Author contributions

LS, CL, AO and AM conceptualized the study and its methodology. LS and AM collected and analyzed the data. LS and AM wrote the first draft. All authors reviewed and edited the final manuscript.

## Transparency declaration

The authors have nothing to disclose.

